# Identification of arginine phosphorylation in *Mycolicibacterium smegmatis*

**DOI:** 10.1101/2021.04.05.438432

**Authors:** Emmanuel C. Ogbonna, Henry R. Anderson, Karl R. Schmitz

## Abstract

Tuberculosis is a leading cause of worldwide infectious mortality. The prevalence of multidrug-resistant *Mycobacterium tuberculosis* (*Mtb*) infections drives an urgent need to exploit new drug targets. One such target is the ATP-dependent protease ClpC1P1P2, which is strictly essential for viability. However, few proteolytic substrates of mycobacterial ClpC1P1P2 have been identified to date. Recent studies in *Bacillus subtilis* have shown that the orthologous ClpCP protease recognizes proteolytic substrates bearing post-translational arginine phosphorylation. While several lines of evidence suggest that ClpC1P1P2 is similarly capable of recognizing phosphoarginine-bearing proteins, the existence of phosphoarginine modifications in mycobacteria has remained in question. Here, we confirm the presence of post-translational phosphoarginine modifications in *Mycolicibacterium smegmatis* (*Msm*), a nonpathogenic surrogate of *Mtb*. Using a phosphopeptide enrichment workflow coupled with shotgun phosphoproteomics, we identify arginine phosphosites on several functionally diverse targets within the *Msm* proteome. Interestingly, phosphoarginine modifications are not upregulated by heat stress, suggesting divergent roles in mycobacteria and *Bacillus*. Our findings provide new evidence supporting the existence of phosphoarginine-mediated proteolysis by ClpC1P1P2 in mycobacteria and other actinobacterial species.

## Introduction

Tuberculosis is a leading cause of worldwide infectious mortality, ranking above HIV/AIDS and Ebola, and is one of the top ten leading causes of death overall (WHO, 2021). Advances in diagnosis, vaccinations and therapeutics have reduced tuberculosis morbidity and mortality. However, the prevalence of multidrug-resistance in the causative bacterium, *Mycobacterium tuberculosis* (*Mtb*), remains high (WHO, 2021). These statistics underscore the urgent need to discover new drugs and exploit new molecular targets. One promising target is the mycobacterial Clp protease. Several studies show that Clp protease components are strictly essential for *Mtb* viability (Sassetti et al., 2003; Griffin et al., 2011; Raju et al., 2012; Raju et al., 2014; DeJesus et al., 2017) and are viable targets for anti-*Mtb* therapeutics (Compton et al., 2013; Gavrish et al, 2014; Gao et al, 2015; Moreira et al., 2015; Famulla et al., 2016; Moreno-Cinos et al., 2019).

Clp proteases mechanically unfold and destroy native cytosolic proteins (Sauer and Baker; 2011; Baker and Sauer 2012). These large enzymatic complexes consist of a core peptidase and an associated hexameric ATP-dependent unfoldase. The mycobacterial Clp peptidase, ClpP1P2, is a heteromer composed of distinct ClpP1 and ClpP2 rings that stack face-to-face to create a barrel-shaped tetradecamer (Akopian et al., 2012; Schmitz et al., 2014; Schmitz et al., 2014b; Li et al., 2016; Vahidi et al., 2020). ClpC1 is one of two mycobacterial unfoldases (the other is ClpX) that can assemble with ClpP1P2 to form a functional protease. ClpC1 is an 848-residue protein with a globular N-terminal domain (NTD) and two AAA+ (ATPases associated with various cellular activities) modules. Ring-shaped hexamers of ClpC1 dock coaxially on the surface of the ClpP2 face of the peptidase (Leodolter et al., 2015; Nagpal et al., 2019). Protein substrates destined for destruction by ClpC1P1P2 must first be recognized by ClpC1. The unfoldase therefore serves a critical regulatory role in proteolysis.

Clp proteases have become major targets of novel drug development against *Mtb* and other pathogenic bacteria. Several classes of antimicrobials specifically target the Clp peptidase, including dysregulators (e.g., acyldepsipeptides (Brötz-Oesterhelt et al., 2005; Kirstein et al, 2009)), activators (e.g., sclerotiamide (Lavey et al., 2016)), and catalytic inhibitors (e.g., β-lactones and boronate compounds (Compton et al., 2013; Akopian et al., 2015)). Importantly, multiple cyclic peptides have been discovered that exhibit anti-*Mtb* activity through dysregulation of ClpC1, including cyclomarin A, lassomycin, ecumicin, rufomycin and metamarin (Schmitt et al., 2011; Gavrish et al., 2014; Gao et al., 2015; Wolf et al., 2019; Li et al., 2021). These compounds bind to the ClpC1 NTD and disrupt proteolysis by uncoupling unfoldase activity from proteolysis or by causing uncontrolled degradation of cellular proteins. The antimicrobial activity of ClpC1-targeting compounds underscores the importance of ClpC1P1P2 in mycobacteria. However, the specific proteolytic functions responsible for its essentiality remain obscure, and only a handful of protein substrates have been identified (Barik et al., 2010; Gopal et al., 2020; Lunge et al., 2020; Ziemski et al., 2021).

An expanded understanding of the physiological roles played by ClpC1P1P2 would bolster efforts to develop antimicrobial compounds. In other bacteria, Clp proteases participate in cellular processes ranging from targeted pathway regulation to cell-wide protein quality control (Gerth et al., 2008; Trentini et al., 2016). Multiple mechanisms of substrate recognition have been described, including direct interactions with the unfoldase (Gopal et al., 2020; Lunge et al., 2020) and indirect recognition with aid of adaptors (Marsee et al., 2018; Ziemski et al., 2021). Recent studies in *Bacillus subtilis* and related Firmicutes have demonstrated that post-translational arginine phosphorylation marks some proteins for destruction by ClpCP (Schmidt et al., 2014; Trentini et al., 2016; Junker et al., 2018). Dual phosphoarginine (pArg) binding sites on the *B. subtilis* ClpC NTD allow ClpCP to recognize phosphoarginylated proteins as proteolytic substrates (Trentini et al., 2016). Phosphoarginine-mediated proteolysis in these bacteria is upregulated during stress through activation of the arginine kinase McsB (Fuhrmann et al., 2009). Phosphoproteomic studies have uncovered widespread arginine phosphorylation during heat stress, implicating ClpCP in turnover of misfolded proteins (Schmidt et al., 2014; Trentini et al., 2016; Trentini et al., 2018). Additionally, McsB regulates the global stress response through targeted phosphorylation of the negative transcriptional regulators CtsR and HrcA (Fuhrmann et al., 2009; Schmidt et al., 2014; Suskiewicz et al., 2019). Proteolysis of these targets by ClpCP allows transcriptional activation of stress response genes (Fuhrmann et al., 2009). Interestingly, ClpC and ClpP are themselves targets of McsB, and specific pArg sites on ClpC are required for its activation by McsB (Schmidt et al., 2014, Elsholz et al., 2012). These studies underline the role of arginine phosphorylation in Firmicutes as both a degradation signal and a regulatory mechanism.

Several lines of evidence suggest that an analogous pArg-mediated proteolytic pathway exists in mycobacteria. Sequence and structural data reveal overall homology between the NTDs of *Mtb* ClpC1 and *B. subtilis* ClpC, as well as strong conservation of the residues surrounding the pArg-binding sites (Trentini et al., 2016; Weinhäupl et al., 2018). Moreover, *in vitro* experiments confirm that the *Mtb* ClpC1 NTD does indeed interact with both free pArg and with arginine-phosphorylated model substrates (Weinhäupl et al., 2018). However, the existence of pArg-mediated proteolysis in mycobacteria remains in question, as no mycobacterial McsB homologs are known and phosphoarginine modifications have not yet been described in these bacteria.

Here, we confirm the existence of post-translational phosphoarginine modifications in *Mycolicibacterium smegmatis* (*Msm*), a nonpathogenic surrogate of *Mtb*. Using a phosphopeptide enrichment workflow coupled to shotgun phosphoproteomics, we identify arginine phosphosites on several protein targets within the *Msm* proteome. Our findings suggest that these modifications are widespread among actinobacterial species.

## Materials and Methods

### ClpC1 Sequence Analysis

Actinobacterial homologs of *Msm* ClpC1 were identified using HMMER (Eddy, 1995) and aligned using Clustal Omega (Sievers et al., 2011). Fragmentary sequences were omitted from analysis. To reduce overrepresentation of similar taxa (e.g., multiple *Mtb* strains), alignments were pruned such that no two sequences had greater than 90% sequence identity. Positional conservation scores were exported from Jalview (Waterhouse et al., 2009) and plotted as a heatmap in Prism (Graphpad). The sequence logo of the NTD region was generated using WebLogo (Crooks et al., 2004).

### Protein Purification and Binding Assays

The nucleotide sequence encoding the NTD of ClpC1 (codons 1 – 147; referred to as ClpC1^NTD^) was amplified from *M. smegmatis* MC^2^155 gDNA (ATCC) and cloned into a pET22b-derived vector (EMD Millipore) in frame with a C-terminal LPETGG sortase recognition sequence (Glasgow et al., 2016) and 6xHis tag. ClpC1^NTD^ was overexpressed in *E. coli* strain ER2566 (NEB) by induction with 0.5 mM isopropyl β-D-1-thiogalactopyranoside at 30°C for 4 h. Cells were harvested by centrifugation, resuspended in Lysis Buffer (25 mM HEPES, 500 mM NaCl, 10 mM imidazole, 10% glycerol, pH 7.5), and lysed by sonication. ClpC1^NTD^ was purified from clarified lysate by Ni-NTA (Marvelgent Biosciences) and anion exchange (Source Q, Cytiva) chromatography. ClpC1^NTD^ was fluorescently labeled via sortase transpeptidation (Glasgow et al., 2016), by 2 h incubation with sortase A (∼1:30 molar ratio) and a 2-fold molar excess of a Gly-Gly-Asn-Lys-(fluorescein-isothiocyanate) peptide (Biomatik) in PBS Buffer (10 mM Na_2_HPO_4_, 1.8 mM KH_2_PO_4_, 137 mM NaCl, 2.7 mM KCl, 10% glycerol, pH 7.4). Excess peptide was removed by size exclusion chromatography (Superdex 75, Cytiva). Purified ClpC1^NTD-FITC^ was concentrated and stored in CPD Buffer (25 mM HEPES, 200 mM NaCl, 10 mM MgCl_2_, 0.1 mM EDTA, pH 7.0). Binding of phosphoarginine (MilliporeSigma) to 0.1 µM purified ClpC1^NTD-FITC^ was assayed in CPD supplemented with 0.05% Tween 20 and 8 mg/ml BSA by microscale thermophoresis using a Monolith NT.115 (NanoTemper) (Jerabek-Willemsen et al., 2014). Thermophoretic data were fit to a quadratic single site binding equation in Prism (Graphpad).

### Cell Culture Conditions

Liquid starter cultures of *M. smegmatis* MC^2^155 were prepared in 20 mL Middlebrook broth base (HiMedia) containing 0.2% v/v glycerol (Fisher Scientific), 0.05% v/v Tween 80, and grown for 48 hours at 37 °C with 250 rpm orbital shaking. Saturated starter cultures were sub-cultured into 900 mL of fresh media at a starting A_600_ ∼0.02, and grown at 37 °C until A_600_ ∼1.0. 500 mL of culture was collected and added to 400 mL of fresh media. Unstressed control samples were further grown at 37 °C, while samples for heat stress were grown at 50 °C for 21 hours. Cells were harvested by centrifugation at 9,000 × g for 20 min at 4 °C, and pellets were resuspended in 5 mL of ice-cold lysis buffer (25 mM HEPES, 200 mM KCl, 10 mM MgCl_2_, 0.1 mM EDTA, pH 7.5) containing 10 mM ATP (Fisher), 200 μL phosphatase inhibitor cocktail (MilliporeSigma), 30 mM sodium pervanadate (Acros Organics), 3 mM NaF and 3 mM sodium pyrophosphate (both from Fisher). Cells were lysed at high pressure using a microfluidizer (Microfluidics). Lysates were clarified at 15,000 × g for 30 min at 4 °C, and supernatant was stored at −80 °C prior to further processing. Total protein content was estimated by Bradford assay (Bio-Rad).

### Filter-Aided Sample Preparation for Mass Spectrometry

Replicate samples of clarified cell lysates (approximately 10 mg total protein) were prepared for mass spectrometry using filter-aided sample preparation (FASP) (Wiśniewski et al., 2009; Schmidt et al., 2014). Reduction of disulfide bonds was achieved by the addition of 200 mM dithiothreitol (DTT; Fisher), followed by incubation at 56 °C for 50 min. Afterwards, samples were diluted in 7 mL of 0.1 M Tris (Gold Biotechnology), pH 7.5 containing 8 M urea and 25 mM 2-iodoacetamide (both from Acros Organics) in 15 mL tubes and incubated in the dark at room temperature for 45 min to carbamidomethylate cysteines. After alkylation, samples were transferred to an Amicon filter (10,000 MWCO, MilliporeSigma), washed twice with 5 mL 0.1 M Tris, pH 7.5, containing 8 M urea by centrifugation at 4,000 × g, then washed twice with 5 mL 0.1 M Tris, pH 7.5, containing 4 M urea, and finally twice with 5 mL 100 mM ammonium bicarbonate (Honeywell). After the last wash step, retentate was reduced to less than 1 mL by spin concentration.

### In-Solution Trypsin Proteolytic Cleavage

MS Grade Trypsin Protease (Pierce) was dissolved to obtain a 20 µg/mL stock in 100 mM ammonium bicarbonate. Trypsin digestion of processed *M. smegmatis* lysates was performed in an Amicon filter unit at an approximately 1500:1 protein:trypsin ratio overnight at 37 °C. Upon digestion, peptides were recovered by centrifugation at 4,000 × g for 10 minutes, followed by the addition of 0.5 M NaCl. Eluted samples were dried under vacuum in a SpeedVac centrifuge (Thermo Fisher).

### TiO_2_ Enrichment of Phosphopeptides

Phosphopeptides were enriched from tryptic peptide samples using Titansphere TiO_2_ beads (5 µm; GL Sciences) in buffer conditions that limited acid hydrolysis of phosphoarginine (Thingholm et al., 2006; Schmidt et al., 2014; Trentini et al., 2016). 5 mg TiO_2_ resin was resuspended in 1 mL Binding Buffer (300 mg/mL lactic acid, 12.5% acetic acid, 60% acetonitrile, 0.2% heptafluorobutyric acid, pH 4 with NH_4_OH). Lyophilized peptide samples were re-dissolved in the TiO_2_ suspension, incubated for 35 min at 20 °C with gentle agitation, then transferred to graphite spin columns (Thermo Fisher). Unbound peptides were removed by a first wash step with 150 µL Binding Buffer and spun at 2000 × g for 1 min, followed by three wash steps using 400 µL of Wash Solution A (200 mg/ml lactic acid, 75% acetonitrile, 2% trifluoroacetic acid, 2% heptafluorobutyric acid), Wash Solution B (200 mg/mL lactic acid, 75% acetonitrile, 10% acetic acid, 0.1% heptafluorobutyric acid, pH 4 with NH_4_OH), Wash Solution C (80% acetonitrile, 10% acetic acid), respectively. The resin was then incubated with 100 µL Elution Solution 1 (1% NH_4_OH, 30 mM ammonium phosphate) and Elution Solution 2 (1.25% NH_4_OH in 50% acetonitrile) for 15 minutes each. The eluate containing phosphopeptides was collected by centrifugation after each incubation. To remove salts, samples were desalted using a HyperSep C18 column (Thermo Scientific). Samples were lyophilized and stored at −80 °C prior to mass spectrometry.

### Liquid Chromatography - Tandem Mass Spectrometry (LC-MS/MS)

Lyophilized and desalted tryptic digests were resuspended in 20 µL 0.5% acetic acid (pH 4.5). An Orbitrap Eclipse MS (Thermo Scientific) coupled with an Ultimate 3000 nano-LC system and a FAIMS Pro Interface (Thermo Scientific) was used for the LC-MS/MS analysis. Peptide samples were first loaded onto a trap column (PepMap C18; 2 cm x 100 μm I.D.) and afterwards separated at a flow rate of 300 nL/min on an analytical column (PepMap C18, 3.0 μm; 10 cm x 75 mm I.D.; Thermo Scientific). A binary buffer system (buffer A, 0.1% formic acid in water; buffer B, 0.1% formic acid in acetonitrile) with a 165-min gradient (1% to 25% buffer B over 125 min; 25% to 32% buffer B in 10 min, then to 95% buffer B over 3 min; back to 1% B in 5 min, and stay equilibration at 1% B for 20 min) was utilized. To achieve field asymmetric ion mobility spectrometry (FAIMS) separation, multiple CVs (−45, −60 and −80) were applied. For all experiments, the survey scans (MS1) were acquired over a mass range of 375-1500 m/z at a resolution of 60,000 in the Orbitrap. Isolation of precursors was done with a width of 1.6 m/z for MS/MS acquisition. They were subsequently fragmented with higher energy collisional dissociation (HCD) using 30% collision energy with a maximum injection time of 100 ms, and collected in Orbitrap at 15,000 resolution. The dynamic exclusion was set to 60 s, and was shared across different FAIMS experiments. LS-MS/MS data was collected in independent biological triplicates.

### Mass Spectrometry Data Analysis

Proteomic analysis was performed in the Proteome Discoverer software suite (version 2.2, Thermo Fisher). Raw data was searched against the *M. smegmatis* (strain ATCC 700084 / MC^2^ 155) UniProt Reference Proteome (Proteome ID UP000000757; 6,602 entries in total) using Sequest HT (University of Washington and Thermo Fisher) (Eng et al., 1994). Iodoacetamide-mediated cysteine carbamidomethylation was set as a static modification, while methionine oxidation and phosphorylation of arginine, serine, threonine, tyrosine and histidine residues were entered as dynamic modifications. Complete trypsinization with a maximum of two missed cleavages was allowed. Precursor mass tolerance was set at 10 ppm, while allowing fragment ions to have a mass deviation of 0.02 Da for the HCD data. Validation of peptide-spectrum matches (PSM) based on q-value was done using Percolator - with target false discovery rates (FDR) of 1% and 5% for stringent and relaxed validation, respectively. The false discovery rate of high confidence protein and peptide identification was at 1%. Localization probability of phosphopeptide hits were analyzed using the PhosphoRS (ptmRS) node of the Proteome Discoverer software (Taus et al., 2011). Only modifications with a PhosphoRS (ptmRS) score ≥75% were selected.

### Gene Ontology Annotation Analysis

Gene Ontology (GO) annotation was performed using the OmicsBox software suite (Conesa et al., 2005). Homologs were identified by BLAST against the SwissProt/UniProt database (Bairoch et al., 1997), with a minimum expectation value 10^−3^. Homolog annotations were compiled from the InterPro database (Hunter et al., 2009). BLAST and InterPro results were used to generate GO terms in terms of biological process. For proteins still unannotated, direct Uniprot BLAST was performed, and tentative assignment of functional group was based on those of obtained homologs.

### Sequence and Structural Analysis of Arginine Phosphosites

For alignment analysis, a 21-mer sequence was obtained containing 10 residues flanking each side of phosphoarginine. All 21-mers were then aligned in BioEdit (Hall, 1999). Structural information, if available, was obtained from Protein Data Bank (PDB) (Berman et al, 2000). Predicted structures were retrieved using neural network-based machine learning model, AlphaFold (version 2, Jumper et al., 2021) and molecular images were prepared in PyMOL (version 2.5.2, Schrödinger). Protein BLAST (Basic Local Alignment Search Tool) of arginine-phosphorylated proteins was performed in the NCBI suite (Gish and States, 1994). Analysis was restricted to the Corynebacterineae suborder. Sequences were aligned using Clustal Omega (Sievers et al., 2011), and conservation and consensus values obtained in Jalview (Waterhouse et al., 2009).

### Data Availability

The mass spectrometry data from this publication have been submitted to the ProteomeXchange Consortium (http://proteomecentral.proteomexchange.org) via the PRIDE partner repository (Perez-Riverol et al., 2019), and assigned the identifier PXD032083.

## Results

### Phosphoarginine Binding Sites in the ClpC1 NTD Are Conserved Across Actinobacteria

ClpC1 is an essential enzyme in *Mtb* and *Msm* (Sassetti et al., 2003; DeJesus et al., 2017), yet few specific cellular roles or proteolytic substrates of the ClpC1P1P2 protease are known. To gain insight into its cellular function, we began by examining ClpC1 sequence conservation across the phylum Actinobacteria by constructing an alignment of 1,195 actinobacterial orthologs (**Fig. 1A**). Like most other type-II Clp unfoldases, ClpC1 possesses a ∼150 amino acid N-terminal domain (NTD), which likely participates in regulation and substrate recognition, but lacks mechanical function or a direct role in catalysis (Schlothauer et al, 2003; Marsee et al., 2018). Surprisingly, the NTD is as well or better conserved than the D1 and D2 AAA+ ATPase rings, suggesting that the NTD has important and conserved functions across Actinobacteria. This may help explain why naturally occurring antibiotics that target and dysregulate ClpC1 have evolved to specifically bind the NTD (Kirstein et al., 2009; Schmitt et al., 2011; Vasudevan et al., 2013; Gavrish et al., 2014; Bürstner et al., 2015; Gao et al., 2015; Jung et al., 2017).

**Figure 1.**
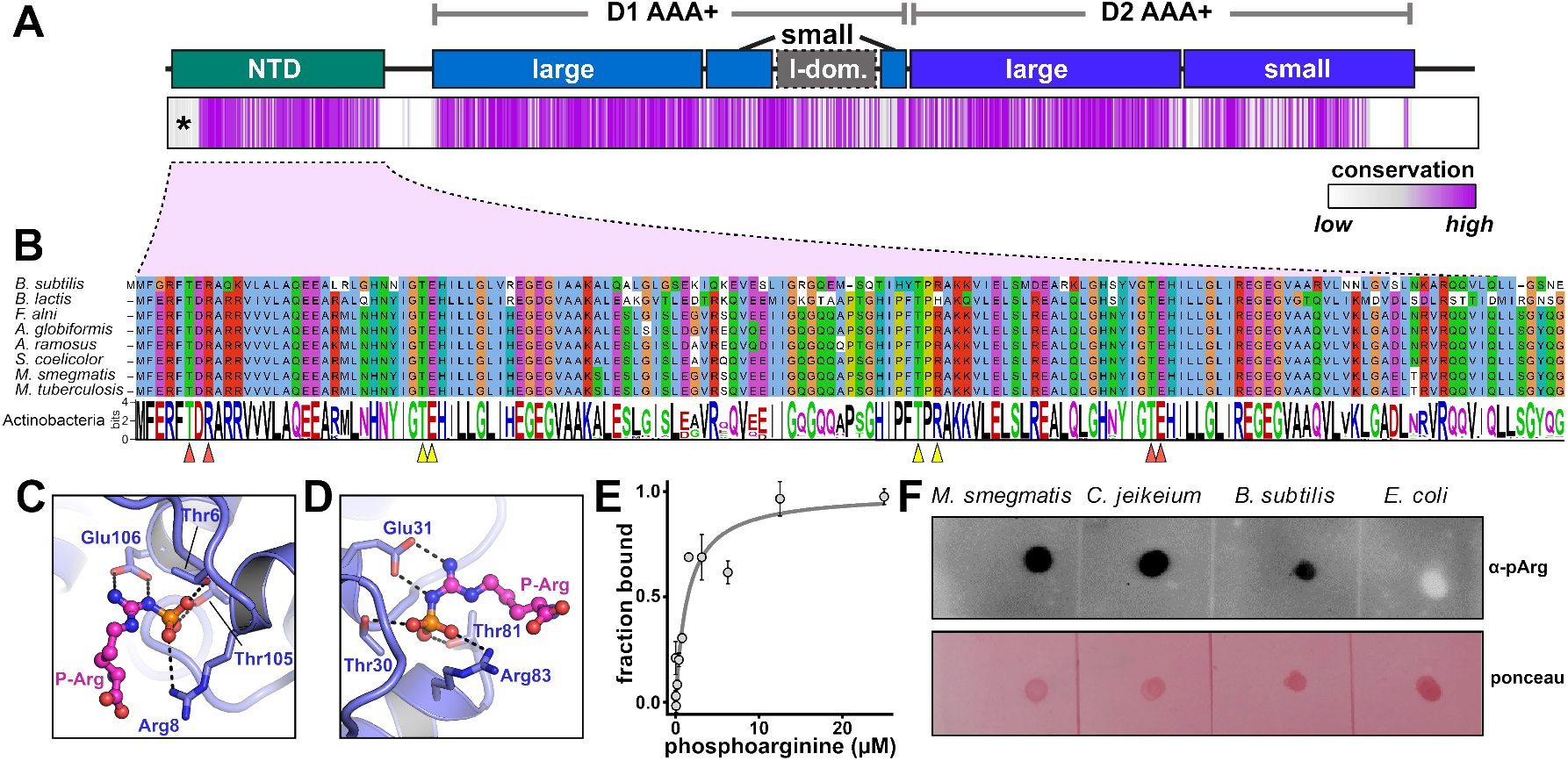
Indirect evidence for phosphoarginine in actinobacteria. **A**) The domain organization of ClpC1 is shown above a plot of amino acid conservation among actinobacterial ClpC1 orthologs, where conservation at each alignment position is plotted as a purple vertical strip. The apparent poor conservation at the beginning of the NTD (marked by *) likely reflects mis-annotation of the start site in some entries. **B**) The sequence of the *Bacillus subtilis* ClpC NTD is shown above equivalent regions from several actinobacterial ClpC orthologs. The sequence logo below shows amino acid conservation in actinobacterial ClpC1 orthologs. Arrows mark positions reported to be important for phosphoarginine binding to site 1 (orange) or site 2 (yellow) (Weinhäupl et al., 2018). Phosphoarginines from the crystal structure of *B. subtilis* ClpC NTD (5HBN) were modeled on the *M. tuberculosis* ClpC1 NTD (6PBQ) putative phosphoarginine-binding sites 1 (**C**) and 2 (**D**) (Wolf et al., 2020). **E**) Binding of phosphoarginine to *Msm* ClpC1^NTD^ was measured by microscale thermophoresis. Data were fit to a noncooperative binding model (gray curve), yielding a *K*_*D*_ of 1.4 ± 0.6 µM. Values are averages of three technical replicates (N = 3) ± one SD. **F**) Dot blots of *Mycolicibacterium smegmatis, Corynebacterium jeikeium, Bacillus subtilis*, and *Escherichia coli* cell lysates, probed with anti-phosphoarginine antibody (Fuhrmann et al., 2015) or stained with Ponceau S.

In *Bacillus subtilis* and other Firmicutes, ClpCP recognizes phosphorylated arginine residues as degradation signals via twin binding sites on opposite ends of the ClpC NTD (Trentini et al., 2016). Examination of actinobacterial ClpC1 sequences revealed that the residues known to be important for phosphoarginine binding in *B. subtilis* are conserved across actinobacterial orthologs (**Fig. 1B**). We found that we could readily model bound phosphoarginines in an existing structure of the *M. tuberculosis* ClpC1 NTD (Wolf et al., 2020), with minor side chains rearrangements, based on the binding mode observed to the *B. subtilis* ClpC NTD (Trentini et al., 2016) (**Fig. 1C,D**). Finally, we directly assessed binding of pArg to purified *Msm* ClpC1^NTD^ by microscale thermophoresis (**Fig. 1E**). Phosphoarginine bound with an affinity of 1.4 µM, similar to the ∼5 µM affinity previously reported to the *Mtb* ClpC1 NTD (Weinhäupl et al., 2018). The ability of the ClpC1^NTD^ to bind pArg and the conservation of pArg binding modules across Actinobacteria provides strong indirect evidence for the existence of this post-translational modification in this phylum.

### Identification of Phosphoarginine Modifications in Mycolicibacterium smegmatis

To test for the physiological existence of arginine phosphorylation, we probed dot blots of several bacterial lysates with a phosphoarginine-specific antibody (**Fig. 1F**) (Fuhrmann et al., 2015). In line with prior studies, signal was detected in *B. subtilis* lysate (Fuhrmann et al., 2015), but not in lysate from *E. coli*, which lacks arginine phosphorylation (Potel et al., 2018, Fu et al., 2020). We also observed a positive immunoblot reaction in lysates from *Msm* and a second actinobacterium, *Corynebacterium jeikeium*. These observations suggest that pArg modifications occur at least within the suborder Corynebacterineae, which encompasses *Corynebacterium, Mycolicibacterium*, and *Mycobacterium*.

To determine which specific cellular proteins carry pArg modifications in *Msm*, we employed an unbiased shotgun proteomics approach. Phosphoarginine is acid-labile and has a short half-life in the acidic TFA-containing solvent systems typically used for proteomic sample preparation and LS-MS/MS (Schmidt et al., 2014). Optimized protocols for arginine phosphoproteomics have been reported that utilize alternative solvent systems at pH ≥ 4 for most steps (Schmidt et al., 2014; Trentini et al., 2016; Trentini et al., 2018). We adapted these methods to our workflow for sample preparation, phosphopeptide enrichment, and LC-MS/MS, thereby minimizing phosphoarginine hydrolysis (**Fig. 2A**). Since stress conditions such as heat shock upregulated the occurrence of this modification in *B. subtilis* (Schmidt et al., 2014), we analyzed lysates from *Msm* cultures grown at either normal growth temperature (37 °C) or 50 °C for 21 hours.

**Figure 2.**
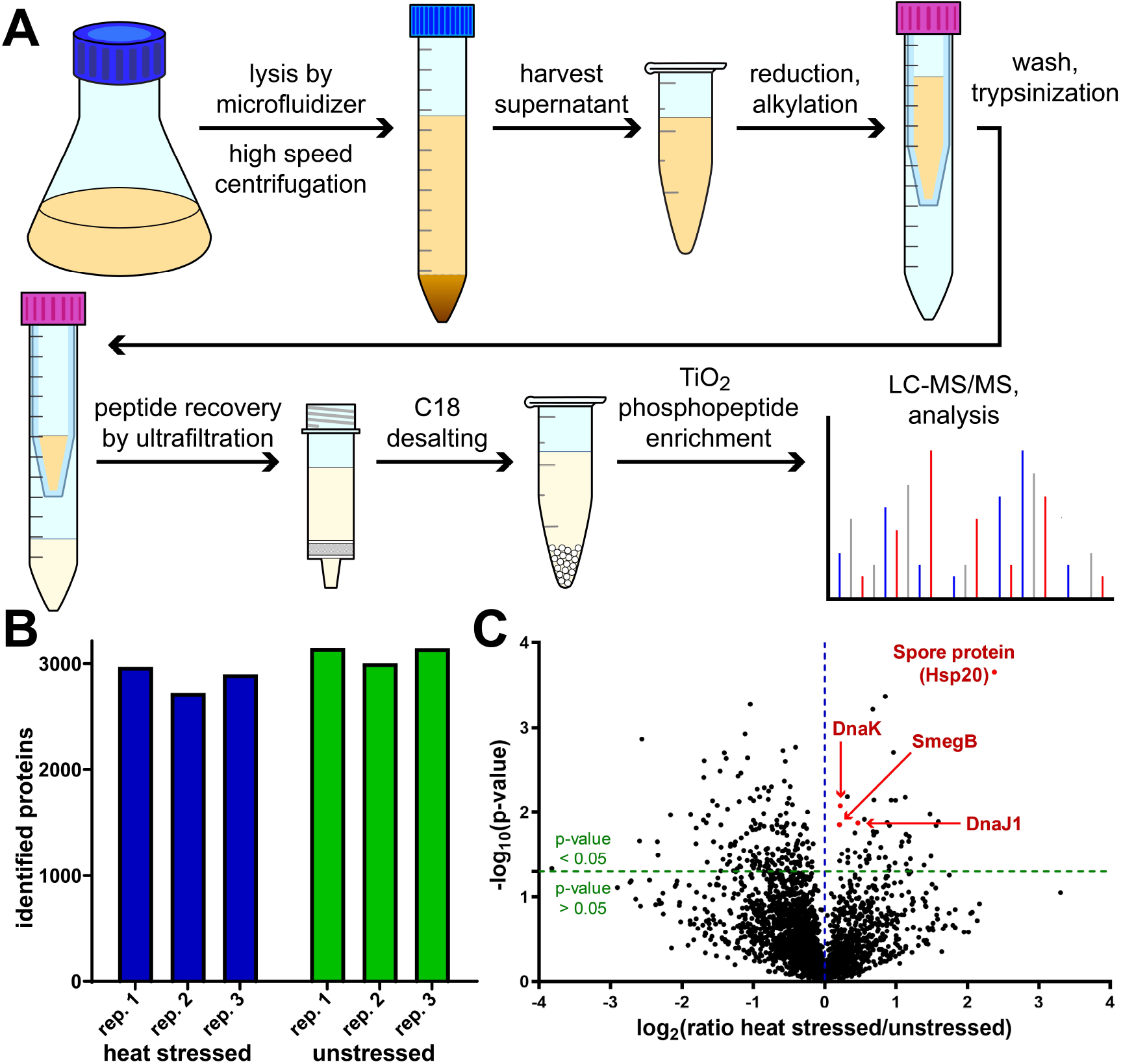
Comparative analysis of mass spectrometry output. **A**) Workflow for arginine phosphoproteomics in *Mycolicibacterium smegmatis*. **B**) Number of proteins obtained across biological replicates from cells grown in heat stress (50°C) and normal conditions (37°C). **C**) Volcano plot showing enrichment of proteins observed by LC-MS/MS in heat stress versus normal growth conditions. Plotted on the *x*-axis is the log^2^ of the ratio of average Sequest HT score of stressed to unstressed samples. The *y*-axis shows -log_10_ of p-values obtained by Student’s t-test. The horizontal green line indicates a cut-off p-value = 0.05; the vertical blue line indicates score ratio = 1. Highlighted in red are notable chaperone proteins with stressed:unstressed ratio > 1.

A similar number (∼3000) of proteins were observed in samples from normal growth (3147, 3004, and 3145 proteins in the three replicates, respectively) and heat stress conditions (2969, 2724 and 2899 proteins) (**Fig. 2B**). This suggests that at this level of stress, compensatory stress responses do not involve a complete shutdown of translational machinery or dramatic upregulation of proteolytic activity. Typically, a major part of stress response to heat shock is the increased expression of heat shock proteins or chaperones. We evaluated the differential levels of observed proteins based on Sequest HT scores. As shown in **Fig. 2C**, several proteins were observed at significantly higher levels (*p*-value ≤ 0.05) in heat shock than in normal growth conditions. These included Spore protein (Msmeg_5611), which belongs to the Hsp20 small heat shock protein family and was enriched over 2-fold upon heat shock. Other chaperone proteins enriched in heat stressed samples included DnaJ1, DnaK, and the SecB-like chaperone SmegB. We also performed comparative GO annotation analysis of strongly enriched/depleted proteins, whose levels differed significantly and by a magnitude of at least two-fold between conditions (**Supplemental Figure S1; Supplemental Table S1**). Proteins involved in amino acid, lipid and noncanonical metabolism, along with redox proteins, were strongly depleted during heat stress. Spore protein was the only stress response protein strongly enriched under heat shock; no protein involved in stress response was strongly depleted. Taken together, this differential expression suggests the activation of canonical heat shock response in these samples.

Using Proteome Discoverer we identified arginine-phosphorylated sites in proteins from these samples. Only peptides with a phosphosite localization probability (PhosphoRS/ptmRS score) ≥75% were selected (Bäsell et al., 2014). We unambiguously localized six phosphoarginine sites in six different proteins (**Fig. 3A; Supplemental Table S2; Supplemental Figure S2**). Most pArg sites were observed in multiple replicates, in both stressed and unstressed samples (**Fig. 3A**). The existence of phosphoarginine modifications in our samples implies the existence of an unidentified *Msm* pArg kinase.

**Figure 3.**
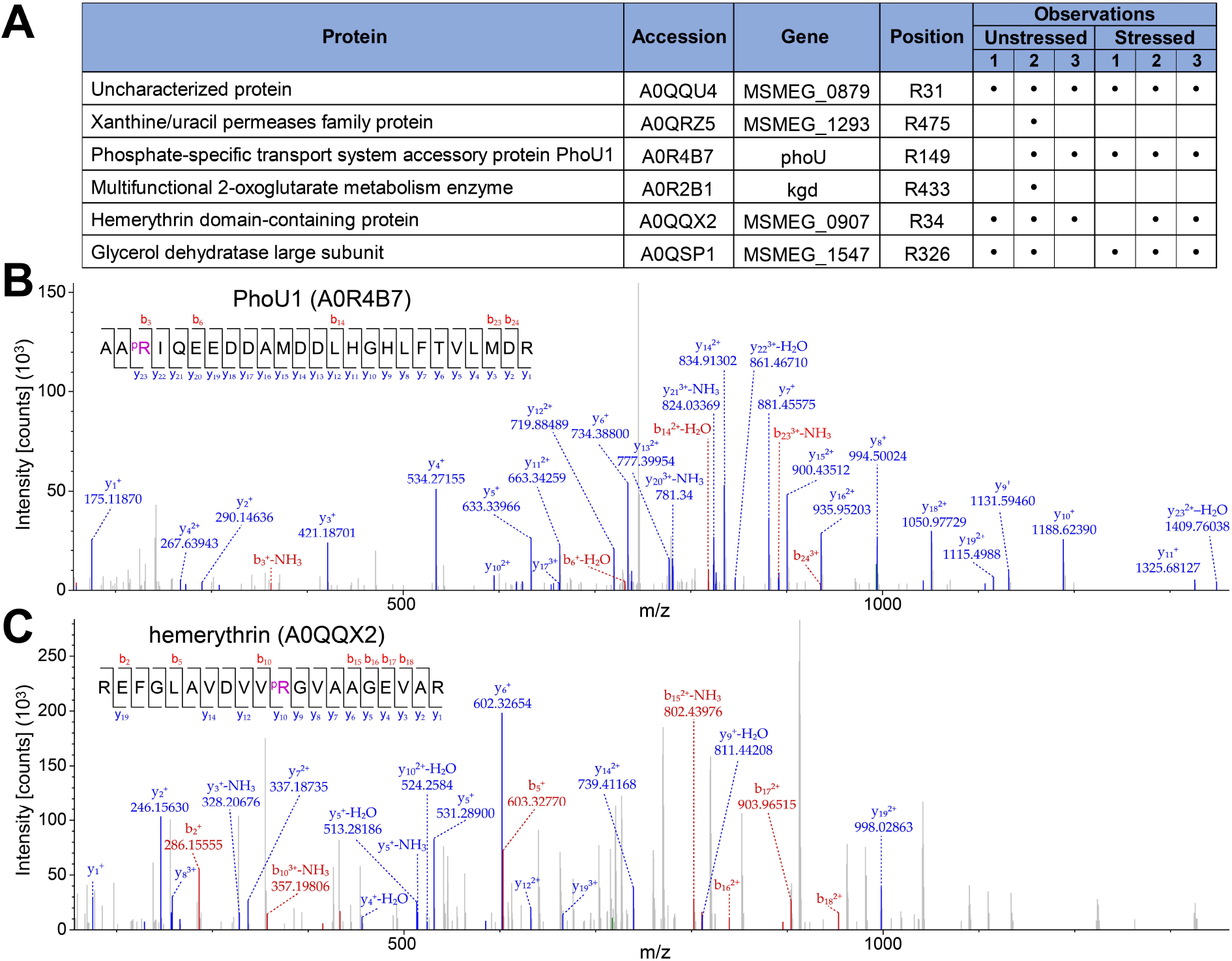
Identification of arginine phosphosites in *Msm* proteome. (**A**) Localized arginine phosphosites in six *Msm* proteins. Localization probability is reported as PhosphoRS/ptmRS score. Representative secondary fragmentation spectra show arginine-phosphorylated peptides from (**B**) Phou1 and (**C**) hemerythrin domain-containing protein. Additional spectra are shown in **Supplemental Figure S2**.

### Functional Characterization of Arginine-Phosphorylated Proteins

We examined the known or predicted functions of the six proteins observed to carry pArg modifications and found that they have diverse functions with no obvious common physiological role (**Fig. 3A**). PhoU1 is a central regulator of the SenX3-RegX3 two-component system responsible for uptake of inorganic phosphate (P_i_) during phosphate starvation (Brokaw et al., 2017). This raises the possibility that pArg modifications are linked to P_i_ availability. Two other pArg-bearing targets were metabolic enzymes: the multifunctional 2-oxoglutarate metabolism enzyme Kgd, involved in the tricarboxylic acid cycle (Wagner et al., 2011), and the glycerol dehydratase large subunit (Msmeg_1547), which potentially contributes to the catabolic pathway involving the glycerol dehydration reaction which yields 3-hydroxypropanal in the presence of adenosylcobalamin coenzyme (**Fig. 3A; Supplemental Figure S2**) (Liu et al., 2010; Nasir et al., 2020). pArg may thus play a role in regulating cellular metabolic pathways. Another phosphosite was detected in the transmembrane xanthine/uracil permease (Msmeg_1293), a nucleobase transporter (**Figs. 3A and 3C**). A site was found on the hemerythrin domain-containing protein Msmeg_0907, which belongs to a class of O_2_-binding proteins involved in signal transduction, response to H_2_O_2_, oxygen sensing and nitric oxide reduction (Isaza et al., 2006; Li et al. 2015; Lo et al., 2016; Ma et al., 2020). Finally, a site was observed on Msmeg_0879, a small (48 aa) uncharacterized protein predicted to be predominantly disordered (**Figs. 3A and B**). Homologs of Msmeg_0879 were detected in close relatives of *Msm*, but are not widely conserved in actinobacteria.

### Structural and Physicochemical Analysis of Arginine Phosphorylation Sites

We assessed whether arginine phosphorylation occurred at sites with particular physicochemical properties. We aligned 21-residue sequence segments centered on each unique phosphoarginine site (**Fig. 4A**) but observe no clear consensus motif. Ala, Leu, Val and Gly appear to be common in flanking positions, although we note that these are the four most abundant amino acids in the *M. smegmatis* proteome (13%, 10%, 9%, 9% of total, respectively) (**Supplemental Table S3**). The sequence diversity surrounding pArg positions suggests that target discrimination is guided by characteristics other than primary sequence.

**Figure 4.**
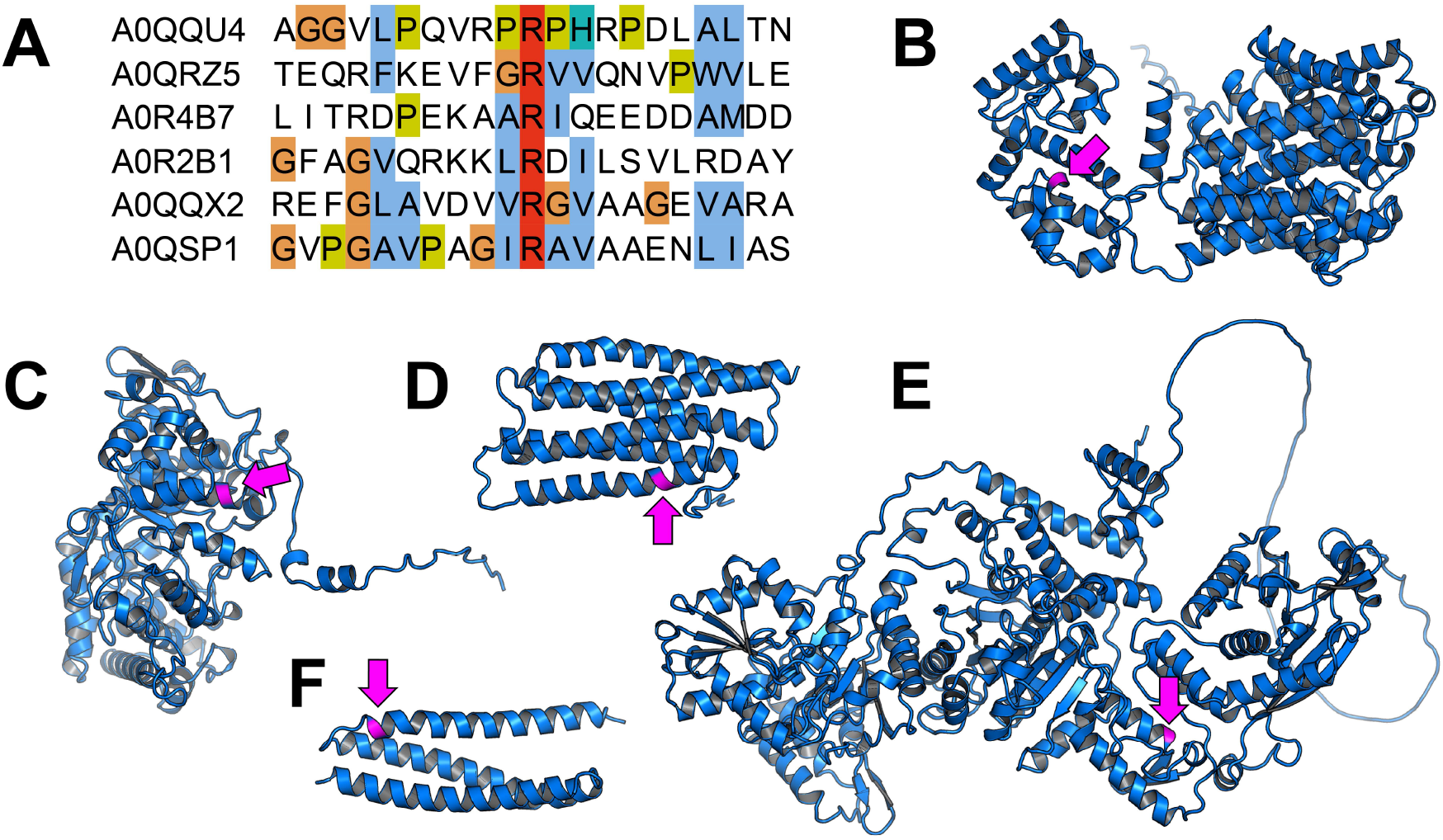
Structural characteristics of phosphoarginine sites. **(A)** Sequence alignment of 20 residues flanking the arginine phosphosites. Structural models of (**B**) Msmeg_1293, (**C**) Msmeg_1547, (**D**) PhoU1, (**E**) Kgd and (**F**) Msmeg_0907 are shown with the phosphorylated arginine colored magenta and marked with an arrow. All models were generated by AlphaFold2 (Jumper *et al*., 2021).

We next examined structural characteristics of phosphorylated positions, based on predicted structures generated by AlphaFold2 (Jumper et al., 2021), except for the small protein Msmeg_0879 for which structure prediction failed. (Notably, the AlphaFold2 prediction for kgd was virtually identical to its reported X-ray structure (PDB ID: 2XT6) with RMSD ∼0.5 A in the region of the phosphosite. (Wagner et al., 2011)) As expected for this charged residue, arginine phosphosites were solvent exposed (**Supplemental Figure S3**), but were located on structured elements rather than loops. Interestingly, in all structures, arginine phosphosites occurred proximal to the beginning or the end of an alpha helix (**Figs. 4B-F**), which may reflect a structural constraint important for recognition by an arginine kinase.

Finally, we assessed the sequence conservation of arginine at these positions. We aligned the arginine-phosphorylated *Msm* proteins with homologs within the Corynebacterineae suborder, and analyzed the positional conservation of arginine at the respective positions. As shown in **Fig. 5**, conservation varies. The phosphosite arginine was well conserved in Kgd (99.8%), Msmeg_0907 (79%), and Msmeg_0879 (62%) (**Figs 5A-C**). Conservation was lower in Msmeg_1547 (34.1%) and PhoU1 (32.63%) (**Figs. 5D and 5E**). Arg was rarely present in this position in homologs of Msmeg_1293 (0.23%) (**Fig. 5F**). In cases where the phosphorylated arginine was well conserved, it may indicate that phosphorylation at this position is a conserved phenomenon with functional or regulatory significance.

**Figure 5.**
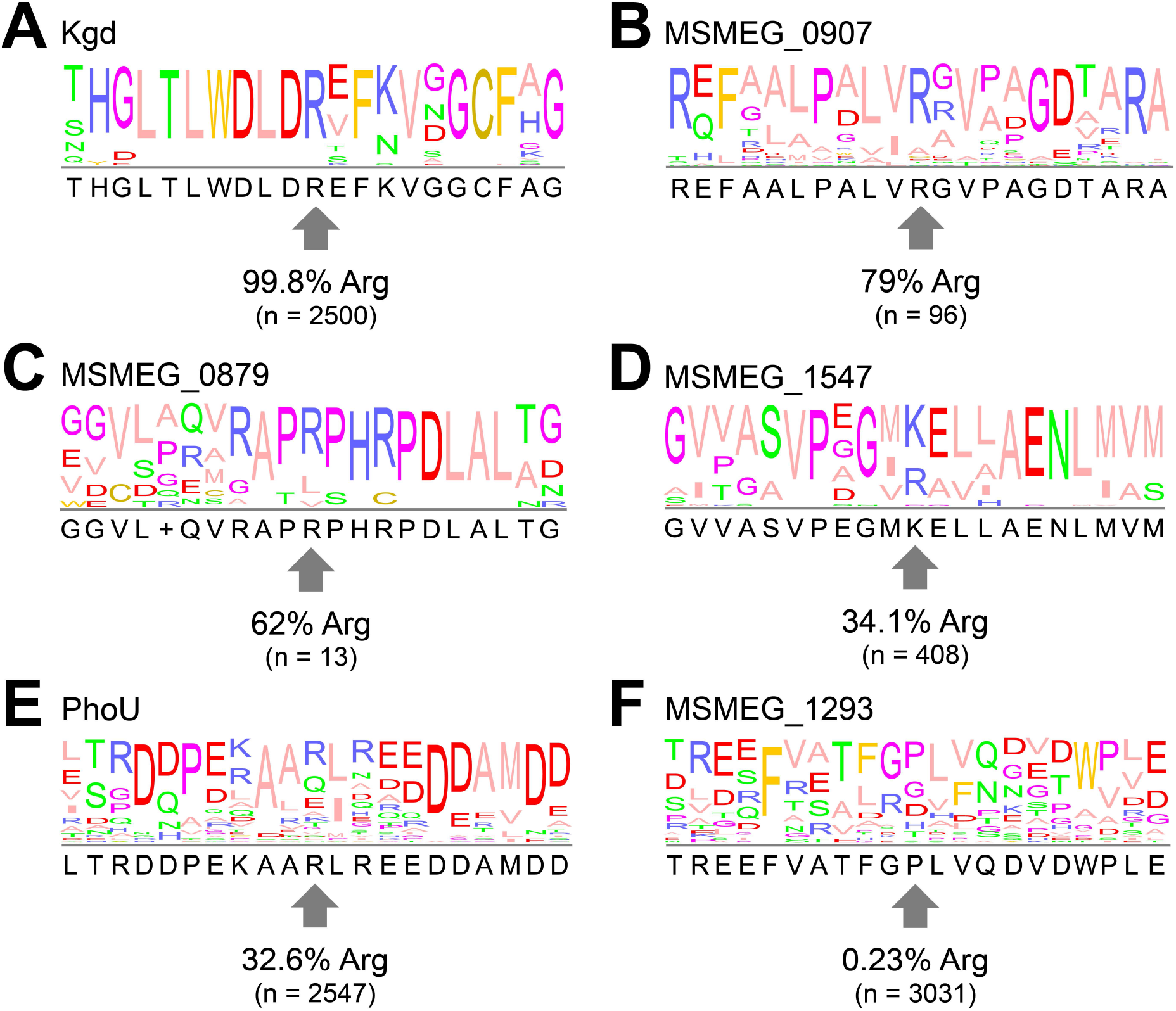
Sequence conservation of phosphorylated arginine residues. Sequence logos show conservation near phosphorylated arginines in (A) Kgd R433, (B) Msmeg_0907 R34, (C) Msmeg_0879 R31, (D) Msmeg_1547 R326, (E) PhoU1 R149, (F) Msmeg_1293 R475. Percentages indicates positional conservation of the phosphorylated arginine; “n” indicates the number of aligned orthologous sequences used to generate the sequence logo.

## Discussion

Protein phosphorylation is a ubiquitous mechanism of signal propagation and pathway regulation in bacteria (Macek et al., 2019). While examples of His, Asp, Ser, Thr, and Tyr phosphorylation are widespread (Chow et al., 1994; Canova et al., 2008; Prisic et al., 2010; Nakedi et al., 2015), the importance of arginine phosphorylation has become apparent only recently. A growing body of data links pArg modifications to protein quality control pathways and spore germination in Firmicutes, including *B. subtilis* (Fuhrmann et al., 2009; Elsholz et al., 2012; Schmidt et al., 2014; Fuhrmann et al., 2016; Trentini et al., 2016; Junker et al., 2018; Weinhäupl et al., 2018; Zhou et al., 2019). However, it has remained unclear how prevalent pArg modifications are in other bacterial phyla. Here we provide direct evidence for the existence of arginine phosphorylation in *Mycolicibacterium smegmatis*, an actinobacterium.

In *B. subtilis* and *S. aureus*, this modification is upregulated by stress (Fuhrmann et al., 2009; Elsholz et al., 2012; Schmidt et al., 2014; Fuhrmann et al., 2016; Trentini et al., 2016; Junker et al., 2018; Weinhäupl et al., 2018; Zhou et al., 2019). Hence, we performed our study by examining pArg levels in cells grown under heat stress and normal conditions. While we saw expected increases in levels of some heat shock proteins overall, arginine phosphorylation abundance was not altered by heat shock, as occurs in firmicutes. It is possible that mycobacterial pArg levels are linked to different stress stimuli that were not tested here. Alternatively, pArg may have no relationship to the mycobacterial stress response, and may instead play a targeted regulatory role.

The six arginine phosphosites localized here were far fewer than those identified in other bacterial studies. Two independent studies in *B. subtilis* observed 121 and 217 sites in 87 and 134 proteins, respectively (Elsholz et al., 2012; Schmidt et al., 2014), while in *S. aureus* 207 sites were identified in 126 proteins (Junker et al., 2018). On its face, the small number of pArg modifications found in *Msm* argues against a proteome-wide quality control role, and appears more consistent with targeted regulation of select proteins and processes. However, we note that phosphoproteomic studies in Firmicutes utilized deletion strains lacking known arginine phosphatases (^*Bs*^YwlE and ^*Sa*^PtpB), which elevate levels of pArg-bearing proteins. No arginine phosphatase has yet been identified in *Msm* or *Mtb*, thus our approach utilized wild-type *Msm*. If a mycobacterial arginine phosphatase is eventually identified, it would be interesting to observe the effect of its knockout or knockdown on pArg levels.

For arginine phosphorylation to be useful to the cell, it must presumably be applied selectively through the regulated activity of arginine kinases. Many protein kinases recognize substrates through characteristic sequence motifs surrounding the phosphosite. By contrast, our examination of mycobacterial arginine phosphosites revealed no clearly enriched consensus sequence around pArg. Prior analysis of Arg phosphosites in *B. subtilis* similarly revealed no evident consensus sequence (Schmidt et al., 2014). This degeneracy suggests either that distal interactions guide *Msm* arginine kinases to phosphosites or that other characteristics of target proteins guide substrate selection. Supporting this latter possibility, we noted that all phosphorylated arginine residues (for which structural models could be obtained) were found near one end of an alpha helix, which may indicate a structural feature recognized by an arginine kinase.

What role does arginine phosphorylation play in mycobacterial cells? In *B. subtilis*, pArg tagging helps enforce protein quality control by marking misfolded proteins for destruction by ClpCP (Elsholz et al., 2012; Schmidt et al., 2014; Trentini et al., 2016). Prior studies have demonstrated that mycobacterial ClpC1P1P2 can degrade model substrates *in vitro* bearing pArg modifications (Weinhäupl et al., 2018). The pArg-bearing proteins identified here may therefore be recognized as proteolytic substrates by ClpC1P1P2. Alternatively, pArg may modulate some aspect of the targets’ function or interactions with other proteins. We observed pArg on PhoU1, which helps regulate P_i_ uptake by inhibiting the activity of the SenX3-RegX3 two-component system when P_i_ is readily available (Brokaw et al., 2017). Additionally, the signal transduction protein Msmeg_0907, metabolic proteins (Kgd and Msmeg_1547), a transmembrane xanthine/uracil transport protein (Msmeg_1293) and an uncharacterized protein Msmeg_0879 were found to be phosphorylated. In terms of essentiality, a prior study showed that the *Mtb* homolog of Kgd is essential for viability (Sassetti et al., 2003); while another report showed that PhoU1 is jointly essential with the PhoU2 protein (Msmeg_1605) for *in vitro Msm* growth (Brokaw et al., 2017). More work is required to determine whether pArg modifications regulate these pathways, and whether this modification plays an essential role in mycobacterial physiology.

This study lays the groundwork for future efforts to expound on the roles of phosphoarginine modifications in mycobacteria. The immediate impediment to further understanding this system is the absence of identifiable orthologs of known arginine kinases or phosphatases. Nevertheless, the lack of consensus motif in the identified phosphosites predicts a promiscuous kinase that targets cellular proteins, in a manner independent of a specific primary sequence. Future work will be required to decipher how arginine phosphorylation is regulated, and how it contributes to the physiology of mycobacteria and other actinobacterial species.

## SUPPLEMENTAL DATA

This article contains supplemental data.

## ACKNOWLEDGEMENTS

We thank members of the Schmitz lab for advice; VP, AJL, PCB, MP, AD, PAO and OU for comments on the manuscript; and the Parashar lab for use of instrumentation. For technical help with mass spectrometry, we thank Dr Yanbao Yu and the University of Delaware Mass Spectrometry Facility. Sortase A was a kind gift from S Novo and JM Fox. The anti-pArg antibody was a kind gift from Dr. Paul Thompson, UMass Medical School.

## Funding and Additional Information

KRS was supported by NIH NIGMS Award P20GM104316. The UD Department of Chemistry & Biochemistry Mass Spectrometry core was additionally supported by NIH NIGMS award P30GM110758-02. This content is solely the responsibility of the authors and does not necessarily represent the official views of the National Institutes of Health.

## Author Contributions

E. O. and K. S. designed research; E. O. and H. A. performed research; E. O. and K. S. analyzed data; E. O. and K. S. wrote the paper.

---

**Supplemental Figure 1.**
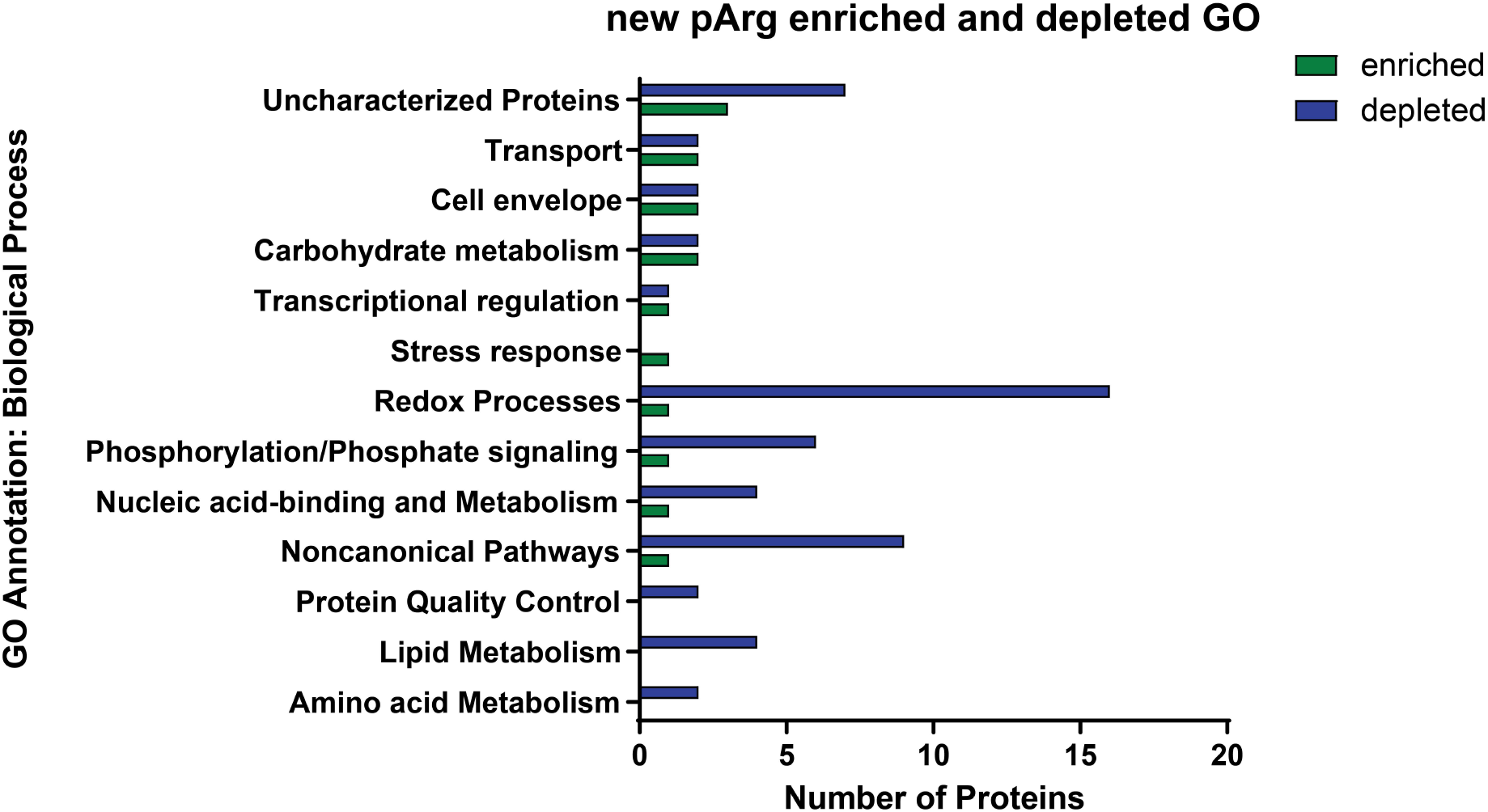
Gene ontology (GO) annotation of proteins strongly enriched or depleted upon heat shock. Proteins with significantly (p ≤ 0.05) and strongly different levels (two-fold or more) in heat shock and control conditions were sorted into GO annotation categories based on biological process.

**Supplemental Figure 2.**
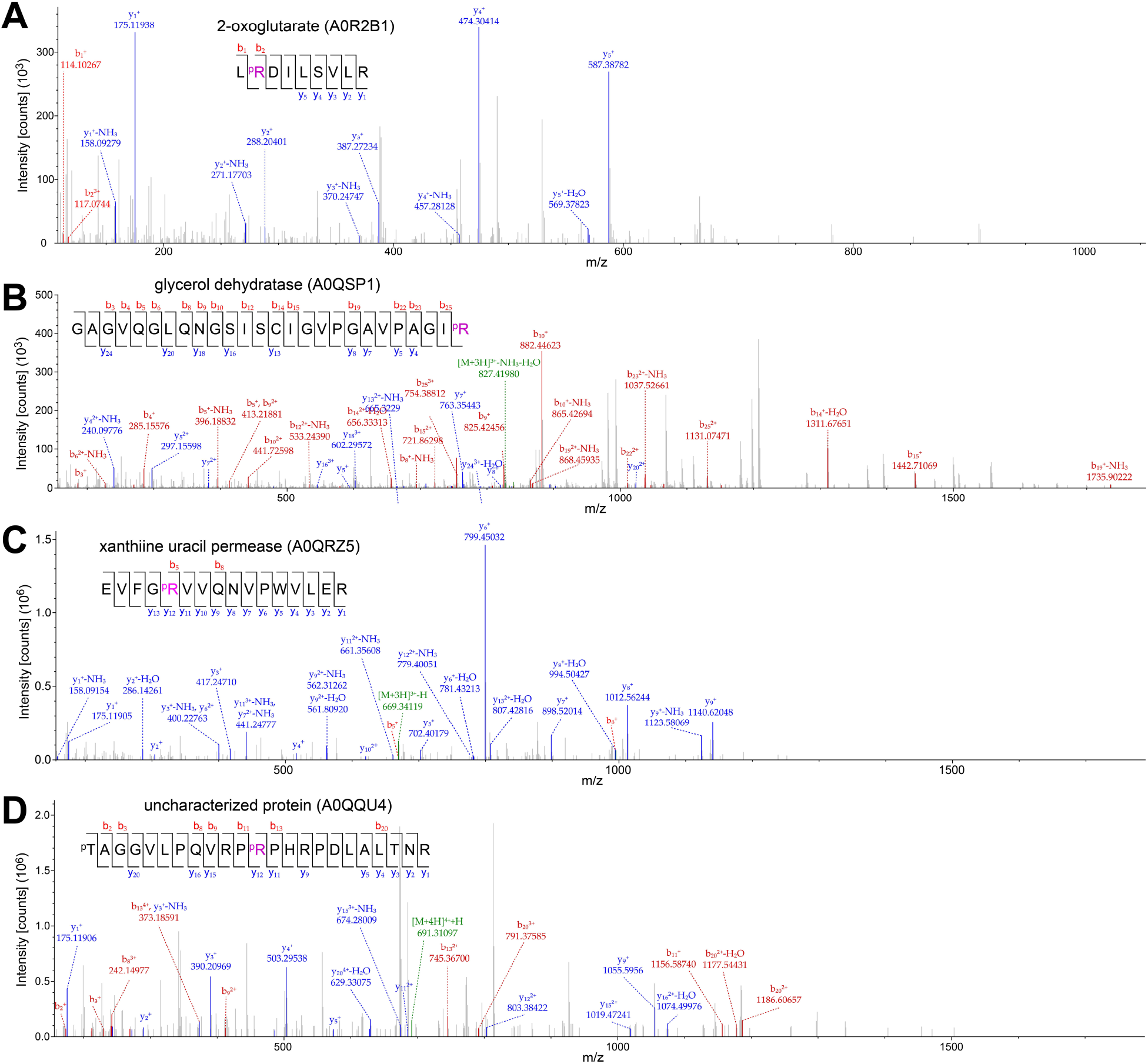
Phosphosite secondary spectra. Secondary fragmentation spectra showing arginine-phosphorylated peptides in (**A**) 2-oxoglutarate, (**B**) glycerol dehydratase, (**C**) xanthine uracyl permease, and (**D**) MSMEG_0879.

**Supplemental Figure 3.**
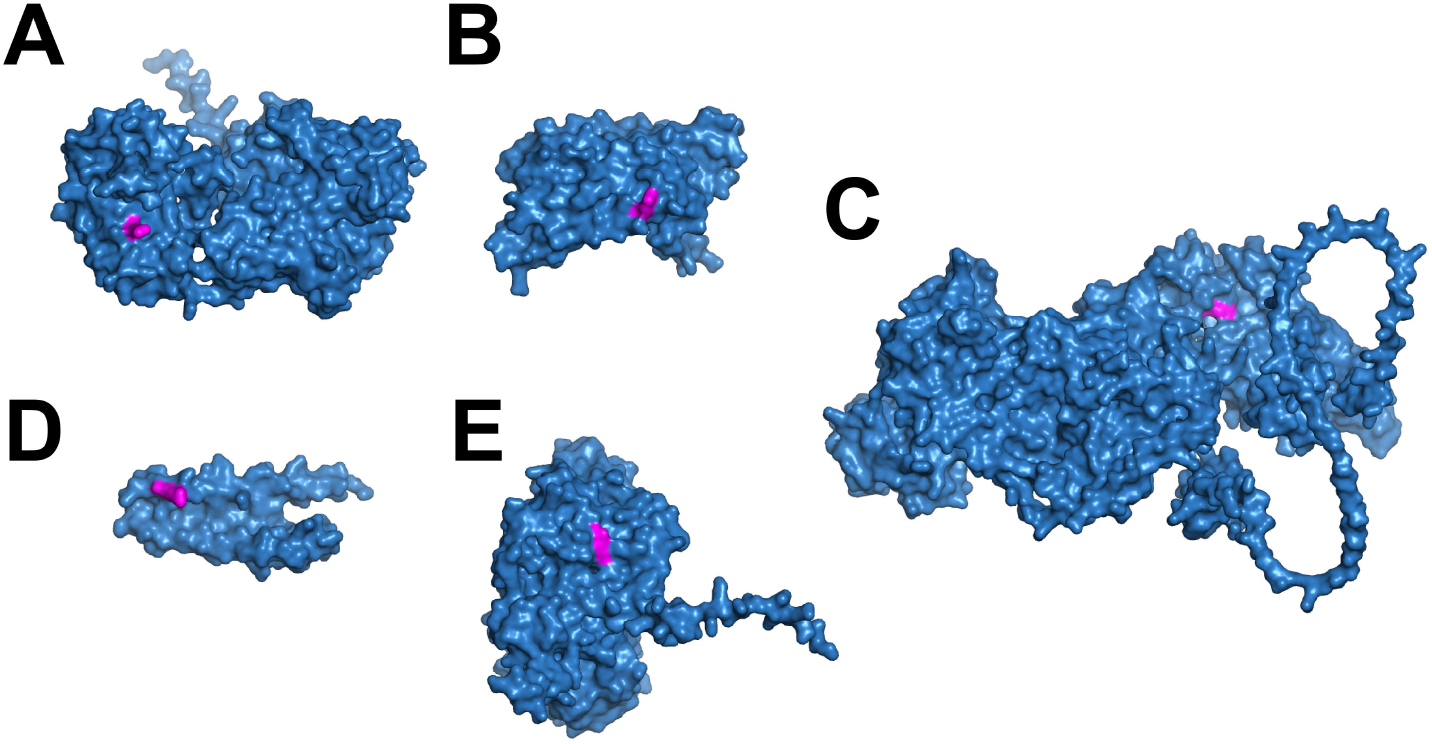
Solvent accessibility of phosphorylated arginines. Surface representations of proteins (**A**) MSMEG_1293 (**B**) PhoU1 (**C**) Kgd (**D**) MSMEG_0907 (**E**) MSMEG_1547 based on AlphaFold2 prediction. Magenta patches represent specific arginine residues observed to be arginine-phosphorylated located on protein surface.

